# Acute exposure to sublethal doses of neonicotinoid insecticides increases heat tolerance in honey bees

**DOI:** 10.1101/2020.10.07.329482

**Authors:** Victor H. Gonzalez, John M. Hranitz, Mercedes B. McGonigle, Rachel E. Manweiler, Deborah R. Smith, John F. Barthell

**Author notes:** Corresponding author, (VHG). These authors contributed equally to this work (Victor H. Gonzalez, John M. Hranitz, Mercedes B. McGonigle, Rachel E. Manweiler). These authors also contributed equally to this work (Deborah R. Smith, John F. Barthell).

## Abstract

The European honey bee, *Apis mellifera* L., is the single most valuable managed pollinator in the world. Poor colony health or unusually high colony losses of managed honey bees result from myriad stressors, which are more harmful in combination. Climate change is expected to accentuate the effects of these stressors, but the physiological and behavioral responses of honey bees to high temperatures while under simultaneous pressure of one or more stressors remains largely unknown. Here we test the hypothesis that exposure to acute, sublethal doses of neonicotinoid insecticides reduce thermal tolerance in honey bees. We administered to bees oral doses of imidacloprid and acetamiprid at 1/5, 1/20, and 1/100 of LD_50_ and measured their heat tolerance 4 h post-feeding, using both dynamic and static protocols. Contrary to our expectations, acute exposure to sublethal doses of both pesticides resulted in higher thermal tolerance and greater survival rates of bees. Bees that ingested the higher doses of pesticides displayed a critical thermal maximum from 2 °C to 4 °C greater than that of the control group, and a reduction in mortality from 69% to 96%. Our study suggests a resilience of honey bees to high temperatures when other stressors are present, which is consistent with studies in other insects. We discuss the implications of these results and hypothesize that this compensatory effect is likely due to induction of heat shock proteins by the insecticides, which provides temporary protection from extremely high temperatures.

## Introduction

Animal pollination is essential for plant reproduction, ecosystem maintenance, and food security, as about 75% of the leading global food crops depend partially or fully on pollinators [1]. The single most valuable pollinator species in the world, and found in both agricultural and natural habitats, is the European honey bee *Apis mellifera* L. [2]. In the U.S. alone, honey bees provide at least $15 billion worth of pollination services and generate from 300 to 500 million dollars in harvestable honey and other products each year [3]. However, managed honey bees are under pressure from myriad stressors that include habitat loss, parasites, diseases, pesticides, and poor nutrition. Bees are now exposed to multiple, simultaneous stressors throughout their lives, which has resulted in unusually high annual colony losses or significant declines in colony health [4].

Several studies demonstrate that the combined effects of stressors are more harmful to bees than one stressor alone [4]. For example, exposure to sublethal doses of neonicotinoid insecticides and nutritional stress renders honey bees more susceptible to the impact of the microsporidian parasite *Nosema* Nägueli, resulting in low brood and adult population [5, 6]. In addition, stressors may act synergistically and thus cause more significant harm to bees. For instance, hive mortality increases when bees are exposed to *Nosema* and to sublethal doses of neonicotinoids or nutritional stress, since the latter two stressors may suppress immunity [7, 8]. Thus, understanding the potential effects resulting from interactions among stressors is relevant for honey bee management and protection.

Climate change is a major new potential stressor altering global temperatures, rainfall, and wind patterns [9]. More severe and frequent extreme weather events are expected, and these will likely accentuate the effects of the stressors that honey bees already face. Alterations in temperature and rainfall are likely to cause spatial, temporal, morphological, and recognition mismatches between plants and pollinators [10]. Changes in the geographical distribution, development, and productivity of honey bees are anticipated, and some studies already document the negative effect of droughts on productivity and survival of honey bee colonies [11]. Climate change will also facilitate the spread of parasites and diseases or intensify their deleterious interactions with honey bees [12]. Clearly, assessing the physiological and behavioral responses of honey bees to high temperatures under the pressure of one or more stressors is a priority.

Few studies have addressed honey bees thermal biology in the presence of other stressors, and the results are not encouraging. For example, acute exposures to diesel exhaust reduces heat tolerance in honey bees, which is concerning because air pollution continues to increase due to rising human population levels and agricultural intensification [13]. Given that insecticides become more toxic at higher temperatures, and that their use is expected to increase under global warming [14], we are therefore interested in determining if insecticides alter the heat tolerance of honey bees.

Here we assess the effect of acute, field-realistic sublethal doses (1/5, 1/20, and 1/100 of LD_50_) of imidacloprid and acetamiprid, two systemic neonicotinoid insecticides that are widely used in agriculture for pest control. Although imidacloprid is more toxic to honey bees than acetamiprid [15], sublethal doses of both insecticides similarly affect their physiology and behavior, such as learning and memory performance, homing ability, foraging, immunocompetence, and susceptibility to parasites [16–19]. To assess the effects of each insecticide on the heat tolerance of honey bees, we use dynamic (ramping) and static (constant) protocols. In the dynamic protocol, we measure bees’ critical thermal maximum (CT_Max_) or the temperature at which an organism loses motor control [20]. Such a physiological parameter has a strong predictive power for understanding bees’ responses to changes in climate as well as land use [21–23]. In the static protocol, we measure bee survival after constant heat exposure. The results of this experiment will be informative about the effects of insecticides on bees’ ability to tolerate a heat stress event. Given the synergistic effects of neonicotinoids with other stressors [7, 8], we hypothesize that bees exposed to acute sublethal doses of insecticides will display a lower CT_Max_ and have a reduced rate of survival in comparison to individuals that are not exposed to insecticides.

## Material and Methods

We conducted experiments during the summer of 2020 using honey bees from an apiary located at the Native Medicinal Plant Research Garden (39°00’37’’N, 95°12’23’’W, 254 m) of the University of Kansas, Lawrence, Kansas, U.S.A. At the apiary, which had four Langstroth hives, we trained bees to forage at a feeder containing a 1.5 M sucrose solution scented with lavender. For all assays, we captured foraging bees between 9:00 and 10:00 h with a glass vial at the feeder, which we then covered with a net mesh (1 mm in diameter). We kept bees inside a cooler (16–19 °C) until we completed fieldwork. Once in the laboratory, we immobilized bees in a refrigerator (3 °C) for 3–5 min and transferred them to 2 mL plastic vials, which had a small opening (2–3 mm in diameter) at one end and a net mesh on the other. Using a micropipette, we fed bees through the vial’s opening or net mesh with 1.5 M sucrose solution to satiation. As in Hranitz et al. [24], we held bees overnight (21–22 h) at room temperature (21–22 °C) before experimentation to ensure all individuals had a similar motivation to feed.

### Insecticide doses

We used commercial formulations with imidacloprid (Macho^®^ 4.0, Agri Star^®^, Albaugh LLC, Ankeny, IA, USA) and acetamiprid (Ortho^®^, flower, fruit & vegetable insect killer, The Scotts Company LLC, Marysville, OH, USA) to prepare stock solutions of each pesticide. We used commercial formulations because we aimed to simulate field conditions by testing the products commonly applied by farmers. We used distilled water to prepare these stock solutions at a concentration of 407 ng/μL for imidacloprid and 500 ng/μL for acetamiprid. We diluted these stock solutions in 1.5 M sucrose to obtain the concentrations of pesticides used in the experiments. We used doses of each pesticide based on the LD_50_ value calculated from acute contact exposure from a topical application, which are 18 ng/bee for imidacloprid and 7100 ng/bee for acetamiprid [15]. We used the following doses for each insecticide: imidacloprid, 3.6 ng/bee (20 % of the LD_50_), 0.9 ng/bee (5 % of the LD_50_), and 0.18 ng/bee (1 % of the LD_50_); acetamiprid, 1420 ng/bee (20 % of the LD_50_), 355 ng/bee (5 % of the LD_50_), and 71 ng/bee (1 % of the LD_50_). Henceforth, the doses 20%, 5%, and 1% are referred as 1/5, 1/20, and 1/100 of LD_50_. As a control, we used 1.5 M sucrose solution without pesticide. We kept all solutions refrigerated and prepared a new stock every week. We administered 10 μL of treatment solutions to bees orally, as previous studies showed that honey bees freely consumed solutions containing up to 40% of imidacloprid [18]. We measured bees’ CT_Max_ and survival after constant heat exposure at 4 h postfeeding, as previous studies indicated that this is the period in which both insecticides have the most effect on honey bees’ behavior (J. Hranitz, per. obs.).

### CT_Max_ assays

To measure CT_Max_, we followed Gonzalez et al. [25] in placing bees individually in sealed glass vials (7.4 ml; 17 × 60 mm) and submerging them horizontally (attached to a metal tray) at approximately 1 cm in depth within a water bath. We used a water bath with a volume of 12 L controlled by a thermostat (18–100 °C; Bellco Sci-Era Hot Shaker, Vineland, New Jersey). We used a dynamic ramping protocol with an initial temperature of 26 °C, which we increased by 1 °C every 2.5 min with an accuracy of ±0.1 °C. To estimate the temperature inside the tubes, we placed an iButton data logger (weight: 3.104 g; DS1923 Hygrochron™; Maxim Integrated, San Jose, California) inside a glass vial and submerged it in the water bath. Thus, we report the temperature inside the tubes not the temperature displayed by the thermostat of the water bath. Pilot experiments indicated that bees held in similar sealed glass vials adjacent to the water bath at room temperature survived through the duration of the CT_Max_ assays. Thus, observed bees’ responses inside sealed vials during our assays were due to changes in temperature, not to oxygen limitation. As an approximation of the CT_Max_, we used the temperature at which bees lost muscular control, spontaneously flipping over onto their dorsa and spasming [20, 26, 27]. We inspected and rotated each vial to determine if the bees had lost muscle control at every Celsius degree until all bees had reached their upper thermal limit.

### Acute heat stress event

To assess whether acute exposure to sublethal doses of pesticides affect the ability of honey bees to tolerate heat stress, we followed Reitmayer et al. [13] in exposing bees to 43 °C inside an incubator and monitored their survival every hour during 5 hours. We conducted this experiment during three consecutive days for each insecticide, collecting and feeding bees with the same doses as indicated above. We placed bees individually inside glass vials and plugged them with a moistened cotton ball (~ 0.2 mL of distilled water per cotton ball) to ensure enough humidity during the experiment.

### Data analyses

We conducted statistical analyses in R [28] and created boxplots and histograms using GraphPad Prism version 7.04 (GraphPad Software, San Diego, CA, USA). We used a Linear Mixed-Effect Model (LMM) to assess effects of insecticide treatments on the CT_Max_. In this model, treatment served as a fixed factor while date of collection as a random factor. We implemented this model using the lme4 package [29] and assessed the significance of fixed effects using a Type II Wald χ2 test with the car package [30]. We used the lsmeans package [31] to conduct multiple pairwise comparisons with Bonferroni adjustment to assess for differences among groups. We used failure-time analyses to assess for differences in bee survival among insecticide treatments.

We implemented a Cox proportional hazard model using the survival package [32], including collection date as a covariate, and conducting post hoc pairwise comparisons with a Log-rank test. To check for the proportional hazard assumption of each Cox model, we tested for independence between time and the corresponding set of scaled Schoenfeld residuals of each variable (treatment and date of collection) using the functions cox.zph in the survival package and ggcoxzph in the survminer package (S1 and S2 Figs; S4 Table).

## Results

The critical thermal maxima (CT_Max_) of honey bee foragers varied among treatments when we exposed them to acute sublethal doses of both insecticide (imidacloprid: Wald χ^2^ = 41.4; acetamiprid: Wald χ^2^ = 31.4, *DF* = 3 and *P* < 0.001 in both cases). Pairwise comparisons with Bonferroni adjustment detected differences in the CT_Max_ between the control group and all other bees treated with imidacloprid. The CT_Max_ of bees was similar among imidacloprid treatments and, on average, from 2.6 °C to 4.2 °C greater than that of the control group. We found a similar pattern in bees fed with acetamiprid, except that the CT_Max_ of bees fed with the lowest dose was similar to that of the control group. Bees fed with the two highest doses (1/20 and 1/5 of LD_50_) displayed a greater CT_Max_, on average, from 2.3 °C to 4.3 °C higher than the control group and bees fed with the lowest dose (see Fig 1a,b; S1 Table).

**Fig. 1.**
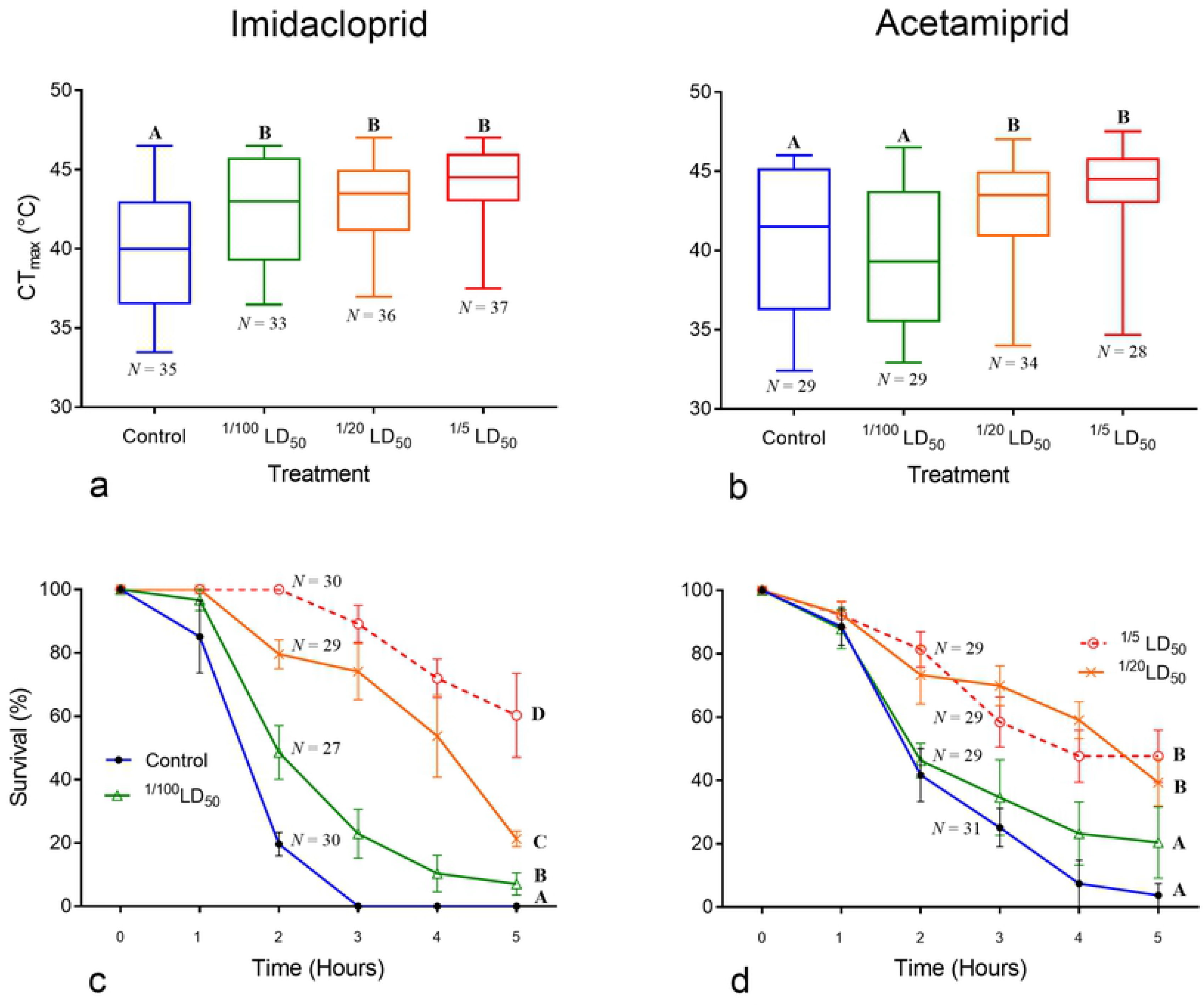
Effect of acute exposure to sublethal doses of neonicotinoid pesticides on the critical thermal maxima (CT_Max_) and survival of honey bee foragers. **a**, **b**, boxplots display average, minimum, and maximum values of CT_Max_. **c**, **d**, bee survival (means ± SE) during a heat stress event (43 °C) over 5 hours. Different letters above boxplots and at the end of each survival curve indicate significant (*P* <0.05) mean differences.

Bee survival also differed among treatments for both insecticide (imidacloprid: Wald χ^2^ = 68.61; acetamiprid: Wald χ^2^ = 25.0, *DF* = 5 and *P* < 0.001 in both cases). In general, survival rapidly decreased over time in bees of both the control group and those fed with the lowest dose (1/100 of LD_50_). Bees fed with higher sublethal doses displayed greater survival rates. In comparison to the control group, hazard ratios (HR) indicated that mortality is reduced from 69% (HR: 0.31) in bees fed with 1/20 LD_50_ of acetamiprid, to 96% (HR: 0.04) in bees fed with 1/5 of LD_50_ of imidacloprid (Table 1). Pairwise comparisons with Bonferroni adjustment indicated differences in the survival of bees among all treatments with imidacloprid (S2 Table). For acetamiprid, bee survival was similar between the control and the lowest dose, each one lower than any of the other two doses (see Fig 1c, d; S3 Table).

**Table 1.**
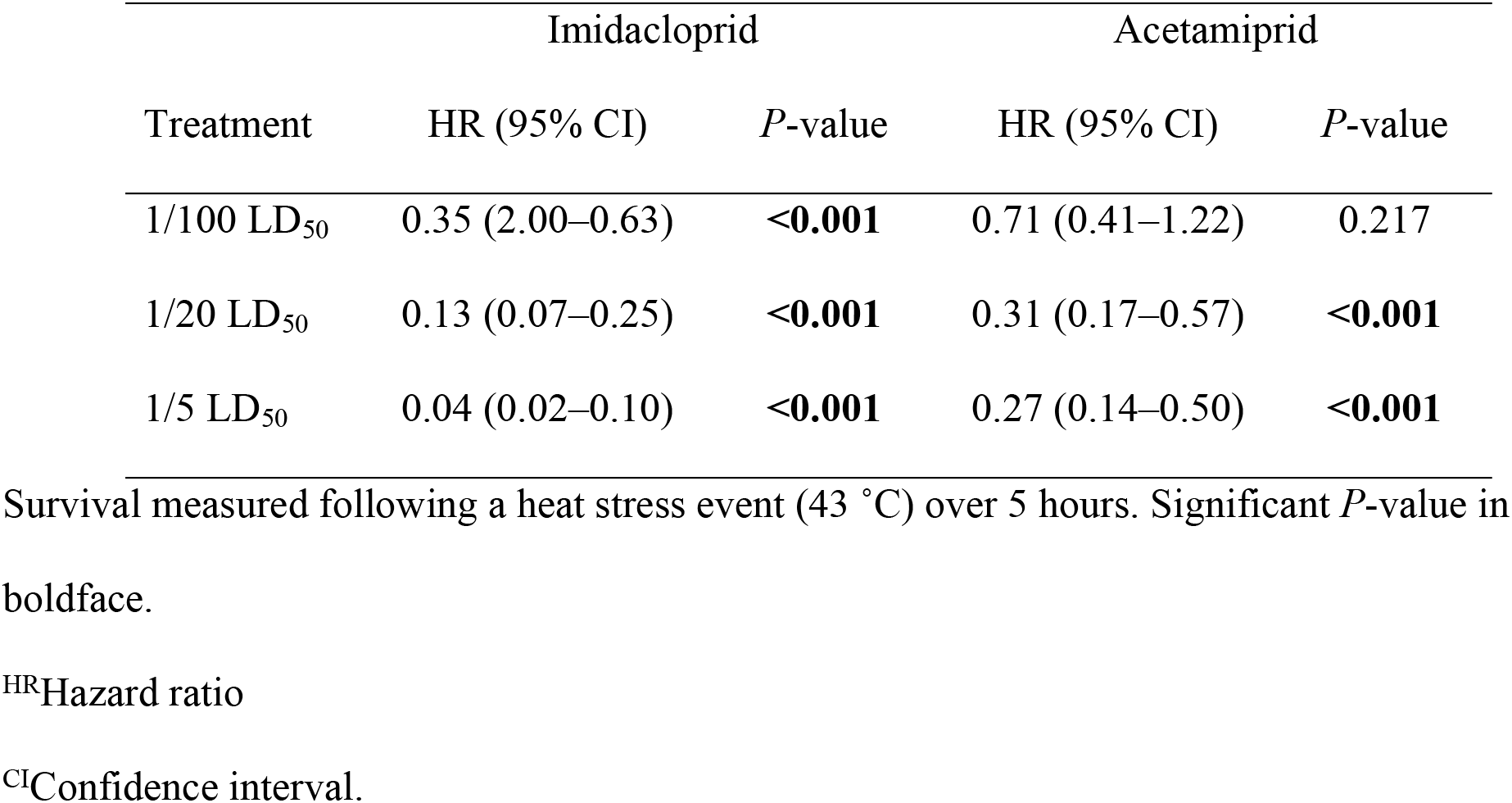
Cox proportional hazards estimates of the survival of honey bees after exposure to three sublethal doses (LD_50_) of imidacloprid and acetamiprid.

## Discussion

The deleterious effects on the development, behavior, physiology, and survival of honey bees due to acute and chronic exposures to lethal and sublethal doses of neonicotinoid insecticides, including imidacloprid and acetamiprid, have been widely documented in the literature [19, 33–36]. Similarly, the synergistic adverse effects of insecticides with other stressors, such as poor nutrition and parasites, have been demonstrated [5–8]. Contrary to our expectations, acute exposure to sublethal doses of imidacloprid and acetamiprid had an positive effect on both honey bees’ CT_Max_ and survival following a heat stress event (43 °C). Bees fed with the higher doses of pesticides (1/20 and 1/5 of LD_50_) displayed a CT_Max_ from 2 °C to 4 °C greater than that of the control group and a reduction in mortality from 69% to 96% (Fig 1, Table 1). Thus, these results do not support the hypothesis that acute, sublethal doses of neonicotinoid insecticides reduce heat tolerance in honey bees.

While unanticipated, our results are consistent with studies in other insect species. For example, Zhang et al. [37] indicate that a pesticide *non-resistant* strain of diamondback moth, *Plutella xylostella* (L.), is more thermotolerant than a resistant strain. As noted by these authors, the greater susceptibility to higher temperatures in the resistant strain likely relates to weaker uploading of heat shock proteins (HSP), among other factors. Heat shock proteins are chaperones that prevent the denaturing of other proteins under heat, as well as under other forms of stress such as cold, starvation, bacterial infections, and exposure to chemicals including pesticides [24, 38]. Inducible heat shock proteins in the HSP70 family of genes are variable in their expression within species, as in the case of the diamondback moth [39]. Similarly, in larval mosquitoes, induced cross-tolerance to a pesticide has been documented through preconditioning at high but sublethal temperatures [40]. Indeed, in honey bees, Koo et al. [41] indicate that heat shock protein expression varies with the type of stressor (including from heat shock), suggesting that pesticides may induce specific responses to various chemical exposures. Thus, we hypothesize that sublethal doses of insecticides activate a stress response in honey bees, which confers further stress resistance to high temperatures. Future studies will attempt to identify this expression profile in correlation with the pesticides used in this work.

The increase in CT_Max_ and greater survival of honey bees after exposing them to sublethal doses of neonicotinoids do not imply any potential benefits to honey bees’ thermal tolerance nor to their resistance to global warming. Instead, our results demonstrate the short-term resilience of honey bees to high temperatures when other stressors are present. The adverse effects on the behavior and physiology of honey bee’s foragers due to neonicotinoid insecticides are unquestionable, including for insecticides with low toxicity, such as acetamiprid, that have been promoted as a “bee-friendly” pesticide in the market. For example, acute chronic sublethal doses of imidacloprid adversely affect aversive learning and reduce overall daily activity, number of foraging trips, and overall lifespan of honey bee foragers [18, 42]. Similarly, sublethal doses of acetamiprid affect locomotor activity, sucrose sensitivity, and memory of honey bees [33]. Thus, although honey bee foragers exposed to acute sublethal doses of insecticides may survive high temperatures, they are behaviorally and physiologically impaired, which in the long-term will alter colony development and productivity.

Acute sublethal doses of pesticides also alter honey bees’ thoracic muscle activity, which allows bees to warm up by shivering their muscles (thermogenesis) and move their wings during flight and fanning the brood. Acute oral exposure to the neonicotinoid thiamethoxam impairs thermogenesis in African honey bees from one hour after exposure and for at least one day, which may not only affect their foraging activity but also other tasks within the colony, such as nest thermoregulation [43]. Similar disruptions to the thermogenic capacity of bees following acute and chronic exposures to both imidacloprid and acetamiprid have been documented in bumble bees [44, 45] and solitary bees [46]. At least under simulated heat wave events, honey bees increase water collection and brood ventilation by recruiting foragers [47]. Because these behaviors require bees to use their thoracic muscles, foragers under the influence of pesticides may be unable to accomplish these tasks successfully, which will influence nest homeostasis.

Honey bee foragers are exposed to pesticides through oral and contact exposures via contaminated nectar, pollen, and/or water. Because of pesticide persistence in the environment, bees are exposed for long periods, not to one, but to a diverse array of pesticides as well as to other agrochemicals that include fungicides and herbicides [48, 49]. However, recent studies demonstrate that exposure to multiple compounds result in additive, synergistic effects, which often increase the toxicity of individual pesticides. For instance, acetamiprid becomes more toxic when combined with triazole fungicides because the latter may inhibit P450-mediated detoxification [49]. Among 98 binary to octonary mixtures of acetamiprid in combination with seven pesticides, 45% of them exhibited synergistic effects on honey bees [50]. Because we used acute sublethal doses of individual pesticides in our laboratory experiments, we do not know if bees would display similar responses to a combination of pesticides and to chronic exposures. It is likely that cumulative toxicity due to a chronic exposure, as well as an increase in toxicity by a combination of pesticides, would inhibit the stress protein response, thus resulting in a lower heat tolerance. Doubtless, future studies should address both factors (combination of agrochemicals and chronic exposures) to obtain a more realistic view of the effects of pesticides on honey bee thermal biology.

Although we tested bees collected from a feeder in order to select foragers, we were unable to control for their age. Several studies have documented a negative relationship between age and heat tolerance in many insects [51, 52], including bumble bees [53]. Thus, the thermal tolerance of honey bees as well as their response to pesticides may vary depending on their age. A mixture of bees from different ages could also explain the high variation in the CT_Max_ observed in our experiments, which ranged from 32 °C to 47 °C across treatments (Fig 1). Second, we measured CT_Max_ as the temperature at which a bee lost muscular control using a dynamic protocol, which requires the visual detection of this physiological event [20]. Detecting this physiological endpoint was particularly challenging in bees that ingested the highest doses of insecticides, which were clearly lethargic from the beginning of the experiment. We are confident with our measurements of CT_Max_ because they are congruent with the results obtained using the static protocol. However, using thermolimit respirometry may be a better approach in these cases, as that method provides a more accurate measurement of CT_Max_ by combining metabolic rate (V_CO2_) and motor activity [54].

To our knowledge, this work is the first in documenting the effects of sublethal doses of pesticides on the heat tolerance of any bee species. Although our results appear counterintuitive at first, they are consistent with results from experiments in other insect species addressing similar questions. As a post hoc hypothesis, we suggest that sublethal doses of insecticides induce the expression of HSPs, which confers further stress resistance to high temperatures. Despite the essential role of temperature and humidity in the development, survival, and health of honey bees colonies [55], as well as concerns about the impact of climate change on pollinators and pollination, it is surprising that the effects of environmental stressors on the bees’ thermal biology have been largely overlooked.

## Acknowledgements

We are indebted to Alex Murray and the University of Kansas’ beekeeping club for allowing us to use the hives, Courtney Goetz, Dieter Schrader, and Robin Lafleur for their assistance with different aspects of the experiments, and Amy Comfort, Mariano Lucia, and anonymous reviewers for comments and suggestions that improved this manuscript.

## Author Contributions

V.H.G. and J.M.H. conceived the experimental design. V.H.G., M.M., and R.M. conducted the experiments. V.H.G. analyzed the data and prepared the first draft. D.R.S. and J.F.B. participated in data analysis and interpretation. All authors contributed in writing the manuscript

## Supporting information captions

**S1 Table. Pairwise comparisons with Bonferroni adjustment of the critical thermal maxima (CT_Max_) displayed by honey bee foragers after acute exposure to sublethal doses of imidacloprid and acetamiprid.** *DF* = 1 in all comparisons. Significant *P*-value in boldface.

**S2 Table. Pairwise comparisons of honey bee survival using a Log-rank test after acute exposure to sublethal doses of imidacloprid followed by a heat stress event (43 °C) over 5 hours.** Significant *P*-value in boldface.

**S3 Table. Pairwise comparisons of honey bee survival using a Log-rank test after acute exposure to sublethal doses of acetamiprid followed by a heat stress event (43 °C) over 5 hours.** Significant *P*-value in boldface.

**S4 Table. Results of test for independence between time and the corresponding set of scaled Schoenfeld residuals of each variable (treatment and date) used in a Cox proportional hazard model.** This model was used to assess for survival of honey bees after exposure to acute sublethal doses of neonicotinoid pesticides (imidacloprid and acetamiprid) followed by a heat stress event (43 °C) over 5 hours. DF = degrees of freedom.

**S1 Figure. Distribution of the scaled Schoenfeld residuals against the transformed time for each variable (treatment and date) of the Cox model built to assess the survival of honey bees after exposure to acute sublethal doses of imidacloprid followed by a heat stress event (43 °C) over 5 hours.**

**S2 Figure. Distribution of the scaled Schoenfeld residuals against the transformed time for each variable (treatment and date) of the Cox model built to assess the survival of honey bees after exposure to acute sublethal doses of acetamiprid followed by a heat stress event (43 °C) over 5 hours.**

## References

1. Klein AM, Vaissière BE, Cane JH, Steffan-Dewenter I, Cunningham SA, Kremen C, et al. Importance of pollinators in changing landscapes for world crops. P Roy Soc B-Biol Sci. 2007; 274: 303–313. https://doi.org/10.1098/rspb.2006.3721

2. Hung K-L J, Kingston JM, Albrecht M, Holway DA, Kohn JR. The worldwide importance of honey bees as pollinators in natural habitats. Proc R Soc B 2018; 285: 20172140.

3. Calderone NW. Insect pollinated crops, insect pollinators and US agriculture: trend analysis of aggregate data for the period 1992–2009. PLoS ONE 2012; 7(5): e37235. https://doi.org/10.1371/journal.pone.0037235

4. Goulson D, Nicholls E, Botías C, Rotheray EL. Bee declines driven by combined stress from parasites, pesticides, and lack of flowers. Science 2015; 347(6229):1255957. doi:10.1126/science.1255957.

5. Wu JY, Smart MD, Anelli CM, Sheppard WS. Honey bees (*Apis mellifera*) reared in brood combs containing high levels of pesticide residues exhibit increased susceptibility to *Nosema* (Microsporidia) infection. J Invertebr Pathol 2012; 109(3): 326–329. doi: 10.1016/j.jip.2012.01.005

6. Branchiccela B, Castelli L, Corona M, Díaz-Cetti S, Invernizzi C, Martínex de la Escalera G, et al. Impact of nutritional stress on the honeybee colony health. Sci Rep 2019; 9: 10156. https://doi.org/10.1038/s41598-019-46453-9

7. Tosi S, Nieh JC, Sgolastra F, Cabbri R, Medrzycki P. Neonicotinoid pesticides and nutritional stress synergistically reduce survival in honey bees. Proc R Soc B 2017; 284: 20171711. https://doi.org/10.1098/rspb.2017.1711

8. Chmiel JA, Daisley BA, Pitek AP, Thompson GJ, Reid G. Understanding the effects of sublethal pesticide exposure on honey bees: a role for probiotics as mediators of environmental stress. Front Ecol Evol 2020; 8(22): 1–19. https://doi.org/10.3389/fevo.2020.00022

9. Seneviratne S, Nicholls N, Easterling D, Goodess C, Kanae S, Kossin J, et al. Changes in climate extremes and their impacts on the natural physical environment. In: Field C, Barros V, Stocker T, Dahe Q, editors. Managing the risks of extreme events and disasters to advance climate change adaptation: special report of the Intergovernmental Panel on Climate Change. Cambridge: Cambridge University Press; 2012. pp. 109–230. doi:10.1017/CBO9781139177245.006

10. Gérard M, Vanderplanck M, Wood T, Michez D. Global warming and plant–pollinator mismatches. Emerg Top Life Sci. 2020; 4(1): 77–86. doi:10.1042/ETLS20190139

11. Flores JM, Gil-Lebrero S, Gámiz V, Rodríguez MI, Ortiz MA, Quiles FJ. Effect of the climate change on honey bee colonies in a temperate Mediterranean zone assessed through remote hive weight monitoring system in conjunction with exhaustive colonies assessment. Sci Total Environ. 2019; 653: 1111–1119. https://doi.org/10.1016/j.scitotenv.2018.11.004

12. Le Conte Y, Navajas M. Climate change: impact on honey bee populations and diseases. Rev Sci Tech Off int Epiz. 2008; 27(2): 499–510. PMID: 18819674.

13. Reitmayer CM, Ryalls JMW, Farthing E, Jackson CW, Girling RD, Newman TA. Acute exposure to diesel exhaust induces central nervous system stress and altered learning and memory in honey bees. Sci Rep. 2019; 9: 5793. https://doi.org/10.1038/s41598-019-41876-w

14. Moe SJ, De Schamphelaere K, Clements WH, Sorensen MT, Van Den Brink PJ, Liess M. Combined and interactive effects of global climate change and toxicants on populations and communities. Environ Toxicol Chem. 2013; 32: 49–61. doi:10.1002/etc.2045

15. Iwasa T, Motoyama N, Ambrose JT, Roe RM. Mechanism for the differential toxicity of neonicotinoid insecticides in the honey bee, *Apis mellifera*. Crop Prot. 2004; 23(5): 371–378. https://doi.org/10.1016/j.cropro.2003.08.018

16. Decourtye A, Devillers J, Cluzeau S, Charreton M, Pham-Delègue M-H. Effects of imidacloprid and deltamethrin on associative learning in honeybees under semi-field and laboratory conditions. Ecotoxicol Environ Saf. 2004; 57: 410–419. doi:10.1016/j.ecoenv.2003.08.001.

17. Yang EC, Chuang YC, Chen YL, Chang LH. Abnormal foraging behavior induced by sublethal dosage of imidacloprid in the honey bee (Hymenoptera: Apidae). J Econ Entomol. 2008; 101: 1743–1748. doi:10.1603/0022-0493-101.6.1743.

18. Karahan A, Çakmak I, Hranitz JM, Karaca I, Wells H. Sublethal imidacloprid effects on honey bee choices when foraging. Ecotoxicology 2015; 24: 2017–2025. https://doi.org/10.1007/s10646-015-1537-2

19. Shi J, Liao C, Wang Z, Zeng Z, Wu X. Effects of sublethal acetamiprid doses on the lifespan and memory-related characteristics of honey (*Apis mellifera*) workers. Apidologie 2019; 50: 553–563. https://doi.org/10.1007/s13592-019-00669-w

20. Lutterschmidt WI, Hutchison VH. The critical thermal maximum: data support the onset of spasms as the definitive end point. Can J Zool. 1997; 75:1553–1560. https://doi.org/10.1139/z97-782

21. Deutsch CA, Tewksbury JJ, Huey RB, Sheldon KS, Ghalambor CK, Haak DC, et al. Impacts of climate warming on terrestrial ectotherms across latitude. Proc Natl Acad Sci USA. 2008; 105: 6668–6672. https://doi.org/10.1073/pnas.0709472105

22. Hamblin AL, Youngsteadt E, López-Uribe MM, Frank SD. Physiological thermal limits predict differential responses of bees to urban heat-Island effects. Biol Lett. 2017; 13: 20170125. https://doi.org/10.1098/rsbl.2017.0125

23. Burdine JD, McCluney KE. Differential sensitivity of bees to urbanization-driven changes in body temperature and water content. Sci Rep. 2019; 9: 1643. https://doi.org/10.1038/s41598-018-38338-0

24. Hranitz JM, Abramson CJ, Carter RP. Ethanol increases HSP70 concentrations in honeybee (*Apis melliera* L.) brain tissue. Alcohol 2010; 44:275–282. https://doi.org/10.1016/j.alcohol.2010.02.003

25. Gonzalez VH, Hranitz JM, Percival CR, Pulley KL, Tapsak ST, Tscheulin T, et al. Thermal tolerance varies with dim-light foraging and elevation in large carpenter bees (Hymenoptera: Apidae: Xylocopini). Ecol Entomol. 2020; 45(3): 688–696. https://doi.org/10.1111/een.12842

26. García-Robledo C, Kuprewicz ER, Staines CL, Erwin TL, Kress WJ. Limited tolerance by insects to high temperatures across tropical elevational gradients and the implications of global warming for extinction. Proc Natl Acad Sci USA. 2016; 113: 680–685. doi: 10.1073/pnas.1507681113.

27. García-Robledo C, Chuquillanqui H, Kuprewicz ER, Escobar-Sarria F. Lower thermal tolerance in nocturnal than diurnal ants: a challenge for nocturnal ectotherms facing global warming. Ecol Entomol. 2018; 43: 162–167. https://doi.org/10.1111/een.12481

28. R Core Team. R: a language and environment for statistical computing. R Foundation for Statistical Computing, Vienna. 2018; available from: https://www.R-project.org

29. Bates D, Mächler M, Bolker B, Walker S. Fitting linear mixed-effects models using lme4. J Stat Softw. 2015; 67(1): 1–48. doi: 10.18637/jss.v067.i01

30. Fox J, Weisberg S. An R companion to applied regression. Second edition. California: SAGE Publications, Inc. 2011

31. Lenth RV. Least-squares means: the R package lsmeans. J Stat Softw. 2016; 69:1–33. doi: 10.18637/jss.v069.i01

32. Therneau T. A package for survival analysis in R. R package version 3.2-3. 2020. Available from: https://CRAN.R-project.org/package=survival.

33. El Hassani AK, Dacher M, Gary V, Lambin M, Gauthier M, Armengaud C. Effects of sublethal doses of acetamiprid and thiamethoxam on the behavior of the honeybee (*Apis mellifera*). Arch Environ Con Tox. 2008; 54: 653–661. https://doi.org/10.1007/s00244-007-9071-8

34. Wu JY, Anelli CM, Sheppard WS. Sub-lethal effects of pesticide residues in brood comb on worker honey bee (*Apis mellifera*) development and longevity. PloS One 2011; 6: e14720. https://doi.org/10.1371/journal.pone.0014720

35. Colin, T, Meikle WG, Wu X, Barron AB. Traces of a neonicotinoid induce precocious foraging and reduce foraging performance in honey bees. Environ Sci Technol. 2019; 53: 8252–8261. https://doi.org/10.1021/acs.est.9b02452

36. Vergara-Amado J, Manzi C, Franco LM, Contecha SC, Marquez SJ, Solano-Iguaran J, et al. Effects of residual doses of neonicotinoid (imidacloprid) on metabolic rate of queen honey bees *Apis mellifera* (Hymenoptera: Apidae). Apidologie 2020; doi: 10.1007/s13592-020-00787-w

37. Zhang LJ, Chen JL, Yang BL, Kong KG, Bourguet D, Wu G. Thermotolerance, oxidative stress, apoptosis, heat-shock proteins and damages to reproductive cells of insecticide-susceptible and -resistant strains of the diamondback moth *Plutella xylostella*. Bull Entomol Res. 2017; 107: 513–526. doi: https://doi.org/10.1017/S0007485317000049

38. González-Tokman D, Córdoba-Aguilar A, Dáttilo W, Lira-Noriega A, Sánchez-Guillén RA, Villalobos F. Insect responses to heat: physiological mechanisms, evolution and ecological implications in a warming world. Biol Rev. 2020; 95(3): 802–821. https://doi.org/10.1111/brv.12588

39. Zhang LJ, Wang KF, Jing YP, Zhuang HM, Wu, G. Indentification of heat shock protein genes hsp70s and hsc70 and their associated mRNA expression under heat stress in insecticide-resistant and susceptible diamondback moth, *Plutella xylostella* (Lepidoptera: Plutellidae). Eur J Entomol. 2015; 112(2): 215–226. DOI: 10.14411/eje.2015.039

40. Patil NS, Lole KS, Deobagkar DN. Adaptive larval thermotolerance and induced cross-tolerance to propoxur insecticide in mosquitoes *Anopheles stephensi* and *Aedes aegypti*. Med Vet Entomol. 1996; 10: 277–282. doi: 10.1111/j.1365-2915.1996.tb00743.x.

41. Koo J, Son T-G, Kim S-Y, Lee K-Y. Differential response of *Apis mellifera* heat shock protein genes to heat shock, flower-thinning formulations, and imidacloprid. J Asia-Pac Entomol. 2015; 18: 583–589. https://doi.org/10.1016/j.aspen.2015.06.011

42. Delkash-Roudsari S, Chicas-Mosier AM, Goldansaz SH, Talebi-Jahromi K, Ashouri A, Abramson CI. Assessment of lethal and sublethal effects of imidacloprid, ethion, and glyphosate on aversive conditioning, motility, and lifespan in honey bees (*Apis mellifera* L.). Ecotoxicol Environ Saf. 2020; 204: 111108 https://doi.org/10.1016/j.ecoenv.2020.111108

43. Tosi S, Démares FJ, Nicolson SW, Medrzycki P, Pirk CWW, Human H. Effects of a neonicotinoid pesticide on thermoregulation of African honey bees (*Apis mellifera scutellata*). J Insect Physiol. 2016; (93–94): 56–63. https://doi.org/10.1016/j.jinsphys.2016.08.010

44. Crall JD, Switzer CM, Oppenheimer RL, Versypt ANF, Dey B, Brown A, et al. Neonicotinoid exposure disrupts bumblebee nest behavior, social networks, and thermoregulation. Science 2018; 362: 683–686. DOI: 10.1126/science.aat1598

45. Potts R, Clarke RM, Oldfield SE, Wood LK, Hempel de Ibarra N, Cresswell JE. The effect of dietary neonicotinoid pesticides on non-flight thermogenesis in worker bumble bees (*Bombus terrestris*) J Insect Physiol. 2018; 104: 33–39. https://doi.org/10.1016/j.jinsphys.2017.11.006

46. Azpiazu C, Bosch J, Viñuela E, Medrzycki P, Teper D, Sgolastra F. Chronic oral exposure to field-realistic pesticide combinations via pollen and nectar: effects on feeding and thermal performance in a solitary bee. Sci Rep. 2019; 9: 13770. https://doi.org/10.1038/s41598-019-50255-4

47. Bordier C, Dechatre H, Suchail S, Peruzzi M, Soubeyrand S, Pioz M, et al. Colony adaptive response to simulated heat waves and consequences at the individual level in honeybees (*Apis mellifera*). Sci Rep. 2017; 7: 3760. https://doi.org/10.1038/s41598-017-03944-x

48. Gill RJ, Ramos-Rodriguez O, Raine NE. Combined pesticide exposure severely affects individual-and colony-level traits in bees. Nature 2012; 491(7422): 105–108. https://doi.org/10.1038/nature11585

49. David A, Botias C, Abdul-Sada A, Nicholls E, Rotheray EL, Hill EM, et al. Widespread contamination of wildflower and bee-collected pollen with complex mixtures of neonicotinoids and fungicides commonly applied to crops. Environ Int. 2016; 88:169–178. https://doi.org/10.1016/j.envint.2015.12.011

50. Wang Y, Zhu YC, Li W. Interaction patterns and combined toxic effects of acetamiprid in combination with seven pesticides on honey bee (*Apis mellifera* L.). Ecotoxicol Environ Saf. 2020; 190: 110100. https://doi.org/10.1016/j.ecoenv.2019.110100

51. Nyamukondiwa C, Terblanche JS. Thermal tolerance in adult Mediterranean and Natal fruit flies (*Ceratitis capitata* and *Ceratitis rosa*): effects of age, gender and feeding status. J Therm Biol 2009; 34: 406–414. https://doi.org/10.1016/j.jtherbio.2009.09.002

52. Chidawanyika F, Nyamukondiwa C, Strathie L, Fischer K. Effects of thermal regimes, starvation and age on heat tolerance of the parthenium beetle *Zygogramma bicolorata* (Coleoptera: Chrysomelidae) following dynamic and static protocols. PLoS ONE 2017; 12: e0169371. https://doi.org/10.1371/journal.pone.0169371

53. Oyen KJ, Dillon ME. Critical thermal limits of bumblebees (*Bombus impatiens*) are marked by stereotypical behaviors and are unchanged by acclimation, age or feeding status. J Exp Biol. 2018; 221: jeb165589. doi:10.1242/jeb.165589

54. DeVries ZC, Kells SA, Appel AG. Estimating the critical thermal maximum (*CT*max) of bed bugs, *Cimex lectularius*: comparing thermolimit respirometry with traditional visual methods. Comp Biochem Physiol A Mol Integr Physiol. 2016; 197: 52–27. doi: 10.1016/j.cbpa.2016.03.003.

55. Abou-Shaara HF, Owayss AA, Ibrahim YY, Basuny NK. A review of impacts of temperature and relative humidity on various activities of honey bees. Insect Soc 2017; 64: 455–463. https://doi.org/10.1007/s00040-017-0573-8

